# AAV-mediated delivery of leptin but not adiponectin improves metabolic health in a mouse model of congenital generalised lipodystrophy

**DOI:** 10.64898/2026.04.07.716869

**Authors:** Nadine Sommer, Ahlima Roumane, Mansi Tiwari, Weiping Han, Lora K. Heisler, George D. Mcilroy, Justin J. Rochford

## Abstract

Lipodystrophies are a group of disorders featuring reduced adipose tissue mass or function, which often leads to significant metabolic disease, reduced lifespan and impaired quality of life. Individuals with congenital generalised lipodystrophy (CGL) have severely reduced adipose tissue mass. The loss of healthy systemic lipid storage typically causes hepatic steatosis and lipoatrophic diabetes. In addition, adipocyte-secreted hormones including leptin and adiponectin are dramatically reduced. Leptin has critical roles regulating appetite and broader effects on lipid and glucose metabolism. Daily injection with recombinant leptin is currently the only specific, approved treatment for CGL. The consequences of adiponectin loss in these patients are not fully understood. Likewise, the potential therapeutic benefit of adiponectin delivery is unclear. Here we examine the effect of delivering leptin or adiponectin by adeno-associated virus (AAV) as potential gene therapy treatment for metabolic disease in CGL using a well-characterised murine model of the condition. AAV-mediated leptin delivery significantly improved hepatic steatosis and hyperinsulinemia. However, adiponectin delivery did not lead to any observed beneficial effects. This demonstrates the potential of gene therapy approaches for long-term delivery of leptin in individuals with lipodystrophy, without the need for continuous supply of perishable therapeutics and painful daily injections.

## INTRODUCTION

Lipodystrophies are a heterogeneous group of rare disorders characterised by reduced or absent adipose tissue and severe metabolic disturbances ^1,2^. Congenital generalised lipodystrophy type 2 (CGL2) is caused by mutations in *BSCL2*, leading to loss of function of the encoded protein seipin and near-complete loss of adipose tissue. Consequently, patients develop profound insulin resistance, early-onset diabetes, hypertriglyceridemia, and hepatic steatosis, frequently progressing to metabolic dysfunction associated steatotic liver disease (MASLD) ^3^. In addition to substantial metabolic disease, lipodystrophy markedly reduces quality of life and life expectancy ^4^. Despite the severity of the condition, therapeutic options remain limited, partly due to its rarity and an incomplete understanding of its pathophysiology^5^.

A hallmark of CGL2 and related lipodystrophies is deficiency of adipocyte-derived hormones, particularly leptin and adiponectin. Circulating leptin levels are markedly reduced, and leptin replacement therapy with metreleptin has demonstrated substantial clinical benefit, improving glycaemic control, hypertriglyceridemia, and hepatic steatosis^6^. However, treatment requires lifelong daily injections, which can be particularly painful due to the lack of subcutaneous adipose tissue in affected individuals. Recombinant leptin is also costly and may be complicated by the development of neutralising antibodies and reduced responsiveness^7^. In rodent models, adiponectin has been shown to ameliorate insulin resistance and fatty liver disease in obesity-associated metabolic dysfunction, yet its therapeutic potential in lipodystrophy remains largely unexplored^8–10^.

Gene therapy could offer an improved way to deliver leptin therapy in CGL, or adiponectin if clinically useful. In contrast to conventional pharmacological treatments, it offers the prospect of long-term efficacy following a single administration. Among available delivery systems, AAV vectors have become a leading platform for *in vivo* applications due to their favourable safety profile and ability to mediate long-term transgene expression in non-dividing tissues^11–13^. Substantial clinical progress has been achieved over the past decade, with multiple AAV-based therapies approved for inherited disorders, highlighting the translational potential of this approach for rare genetic diseases.

In the present study, we investigated whether AAV-mediated gene transfer of leptin or adiponectin could improve metabolic complications in lipodystrophy. Using an established seipin knockout (SKO) mouse model of CGL2, we assessed the impact of long-term adipokine expression on insulin sensitivity and associated metabolic complications, with particular focus on liver steatosis. Based on its high hepatic transduction efficiency, AAV serotype 8 was selected for *in vivo* delivery of human LEPTIN or ADIPOQ^14,15^. Through this approach, we aimed to determine the therapeutic potential of sustained adipokine expression as a gene-therapy based treatment strategy for CGL2.

## MATERIALS AND METHODS

### Animal studies

All SKO mice (C57BL/6J background)^16^ were group housed under standardised conditions with 12-h/12-h light/dark cycle, 20-24 °C, 45-65% humidity and had *ad libitum* access to water and chow diet (CRM (P) 801722, Special Diets Services) unless otherwise stated. Procedures on C57BL/6J and SKO mice were approved by the University of Aberdeen Ethics Review Board and performed under a project license (PPL: PP4951597) approved by the UK Home Office. Tissues from 5h fasted mice were dissected following CO_2_ terminal anaesthesia and cervical dislocation, frozen in liquid nitrogen, and stored at -70 °C for further analysis. A EchoMRI-500 body composition analyser (Zinsser Analytic GmbH) was used to determine body composition. Blood glucose levels were measured by glucometer readings (AlphaTRAK3, Zoetis) from tail punctures.

### AAV administration

Nine-week-old SKO mice were randomised and intraperitoneally (i.p.) injected with 1×10^12^ genome copies of AAV8 expressing either human LEP (NM_000230.3), human ADIPOQ (NM_004797.4) or eGFP (Nucleic Acids Res. 24:4592 (1996)) under the CMV promoter. Wild-type (WT) littermates were used as control mice. Vectors were purchased from VectorBuilder: AAV8-CMV-hLEP: VB220206-1202csh, AAV8-CMV-hADIPOQ: VB220206-1205xjd, AAV8-CMV-eGFP: VB010000-9394npt.

### Oral glucose tolerance test

SKO mice, either i.p injected with AAV8-CMV-eGFP, AAV8-CMV-hLEP or AAV8-CMV-hADIPOQ and WT littermates were fasted for 5h. Mice had *ad libitum* access to water throughout. Basal glucose levels (0 min) were measured by glucometer readings (AlphaTRAK3, Zoetis) from tail punctures before a 2 g/kg D-glucose (Sigma) bolus was given by gavage. Blood glucose levels were monitored at 15-, 30-, 60-, 90- and 120-min as in Roumane *et al*.^17^.

### Insulin tolerance test

SKO mice, either i.p. injected with AAV8-CMV-eGFP, AAV8-CMV-hLEP or AAV8-CMV-hADIPOQ, and WT littermates were fasted for 5h. The mice had access to water *ad libitum*. Basal glucose levels (0 min) were measured by glucometer readings (AlphaTRAK3, Zoetis) from tail punctures before 0.75 IU/kg Insulin (Actrapid 100 IU/mL, Novo Nordisk) was administered by i.p. injection. Blood glucose levels were monitored at 15, 30-60- and 90-min.

### H&E staining

Liver tissue was fixed in 10% formalin and embedded in paraffin. 5 µm sections were cut using a microtome (Leica RM2125RT), deparaffinised and stained with haematoxylin Harris as previously described in Sommer *et al*.^15^.

### Serum analysis

Serum was collected, frozen in liquid nitrogen and stored at -70 °C until analysis. Mouse insulin, human adiponectin, human leptin, and mouse ALT analyses were performed at the Core Biochemical Assay Laboratory (Cambridge, UK).

### Gene expression

Frozen tissue was homogenised in the PRECELLYS®24 system (Bertin Instruments) and total RNA was extracted using the RNeasy Mini Kit (Qiagen). Equal quantities of RNA were treated with DNase I (Sigma) and reverse-transcribed with M-MLV reverse transcriptase (Promega). RT-qPCR was performed using a CFX384 Touch Real-Time PCR Detection System (Bio-Rad). No template and no reverse transcriptase controls were performed for every gene analysed. The geometric mean of three stable reference genes (Nono, Ywhaz, and Hprt) was used for normalisation.

### Liver triglyceride assay

Frozen liver tissue samples were homogenised in 1 mL 1xPBS (Sigma). Liver lysates were centrifuged at 16,000 x g for 5 min at 4 °C. Supernatants were collected and triglyceride (TG) levels determined against a standard curve using Triglyceride Liquid Assay (Sentinel Diagnostics) as in Mcilroy *et al*.^18^ and normalised to individual tissue weight.

### Western blot

Frozen tissue samples were homogenised in RIPA buffer containing protease inhibitors (Roche). After centrifugation, the supernatant was collected, and protein concentration was measured via the Pierce BCA Protein Assay (Thermo Scientific). Equal protein amounts were separated by standard SDS-PAGE (NuPAGE Bis-Tris protein gels, Invitrogen) and transferred to PVDF membranes (Immobilon). Protein expression was detected using following primary antibodies: b-actin (R&D Systems), ADFP/PLIN2 (Abcam), and anti-rabbit or mouse IgG HRP-linked secondary (Cell Signaling). Membranes were visualised using Immobilon Crescendo Western HRP substrate (Millipore) and quantified using ImageJ software (NIH).

### Statistical analyses

Data was analysed using GraphPad Prism 10 and are presented as mean ± SEM. Statistical analyses included unpaired two-tailed Student’s t-test, one-way ANOVA with Tukey’s post-hoc test, two-way ANOVA with Tukey’s post-hoc test or mixed-effects analysis, as appropriate. Normal (gaussian) distribution of the samples was assessed, and statistical analyses were adjusted when normality was not met. A p-value <0.05 was considered statistically significant.

## RESULTS

### AAV delivery of human leptin and adiponectin, and its effect on body composition

To assess the potential of AAV-mediated leptin or adiponectin delivery we designed constructs to generate AAV8 expressing either human leptin or adiponectin under the control of the strong and widely expressed CMV promoter (Fig. 1A, B). A cohort of 8-week-old male SKO mice were subjected to metabolic assessment then i.p. injected with 1×10^12^ genome copies of either AAV-CMV-hLEP (SKO hLeptin), AAV-CMV-hADIPOQ (SKO hAdipoq) or AAV-CMV-eGFP (SKO eGFP) as control. Male WT littermates were included to provide corresponding values for healthy, control mice. Mice were then maintained on a chow fed diet for 7 weeks during which metabolic assessments were performed (schematically represented in Fig. 1C).

**Figure 1.**
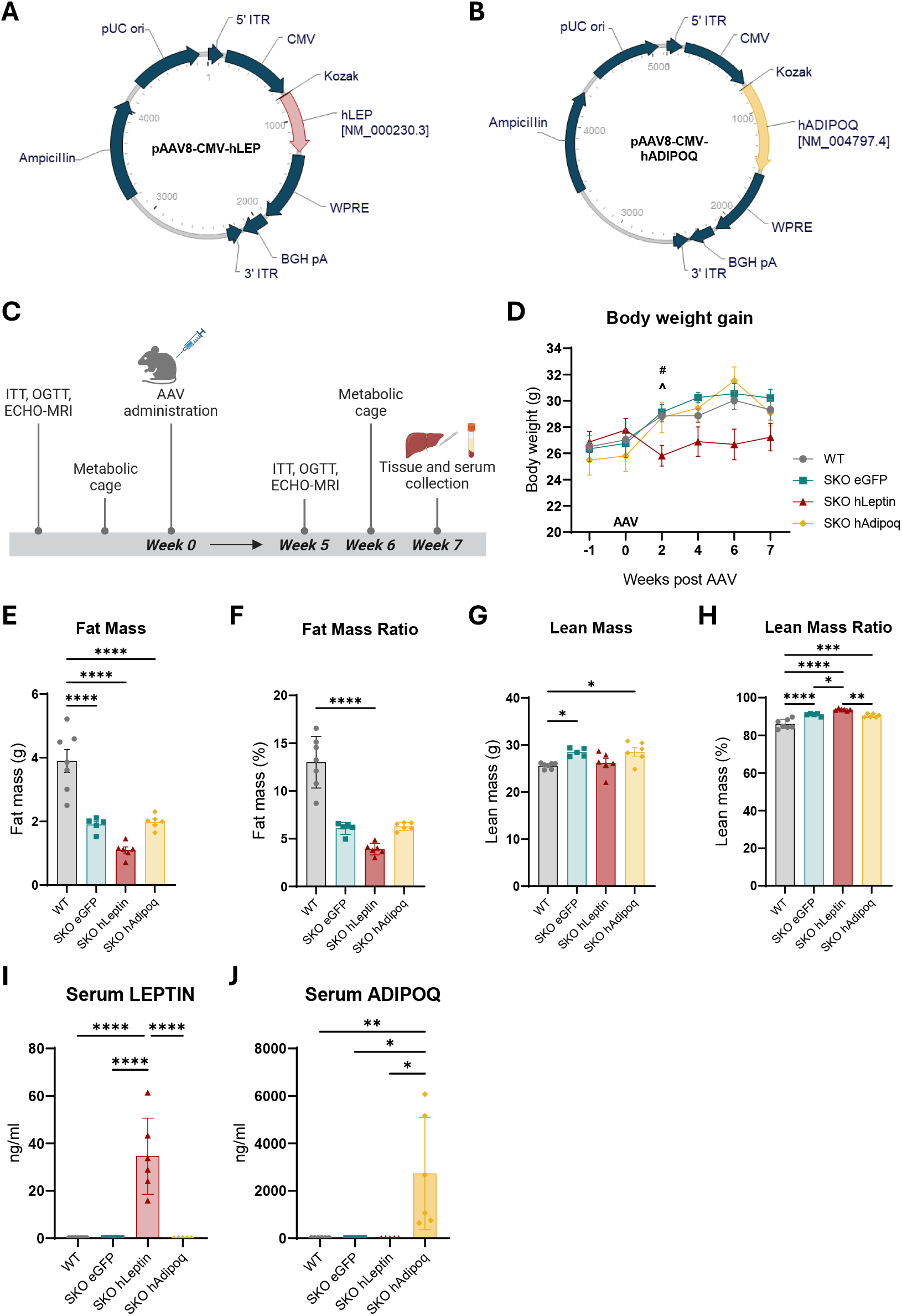
Viral vector plasmid design to overexpress either human ADIPOQ (NM_004797.4) (A) or human LEP (NM_000230.3) (B) transcript driven by the CMV promoter (AAV8-CMV-hADIPOQ or AAV8-CMV-hLEP). Schematic of experimental procedure performed on WT and SKO mice (C), created with BioRender.com. Body weight in WT and SKO mice following gene therapy (D). Whole body fat mass (E), fat mass normalised to body weight (F), lean mass (G) and lean mass normalised to body weight (H) determined by ECHO-MRI 7 weeks post AAV injection. Detection of circulating human leptin (I) and human adiponectin (J) in serum from AAV injected SKO mice fasted for 5 hours. Data presented as the mean ±SEM, for A n = 7 (WT), 8 (SKO eGFP), 6 (SKO hLeptin), 6 (SKO hAdipoq), for D-J n = 7 (WT), 5 (SKO eGFP), 6 (SKO hLeptin), 6 (SKO hAdipoq). For D ^WT - SKO hLeptin, ^#^SKO eGFP – SKO hLeptin. One-way ANOVA/2-way ANOVA, *p≤0.05, **p≤0.01, ***p≤0.001, ****p≤0.0001.

Over the 7-week assessment period SKO eGFP mice gained weight to a similar degree to WT mice (Fig. 1D, Supp. Fig 1A). This is consistent with our previous studies in SKO mice where adipose driven body weight gain in WT mice is approximately matched by increased liver weight gain in SKO mice lacking WAT. Following AAV administration, SKO hAdipoq mice exhibited similar weight gain to the SKO eGFP group (Fig. 1D, Supp. Fig 1A). In contrast, body weight of mice in the SKO hLeptin cohort did not significantly increase during the study period (Fig. 1D, Supp. Fig 1A). Also consistent with our previous studies SKO mice had significantly reduced fat mass and increased lean mass compared with their WT littermates and this was also observed in the SKO hLeptin and SKO hAdipoq mice (Fig. 1E-H).

Analysis of serum revealed robust levels of human leptin and human adiponectin detectable in the SKO hLeptin and SKO hAdipoq mice, respectively. Thus, AAV-mediated delivery of the human transgene resulted in appearance of the adipokines in the circulation (Fig. 1I, J). Because adiponectin therapy represented a novel therapeutic intervention in this disease model we applied the same experimental approach to female mice for WT, SKO eGFP and SKO hAdipoq cohorts, and the corresponding data are presented in Supplementary Figure 2. Females exhibited a comparable pattern to males across all measured parameters, including body composition (Supp. Fig 2A-D) and serum human adiponectin concentrations (Supp. Fig. 2E).

### Leptin but not adiponectin gene therapy improves metabolic complications in SKO mice

To examine if leptin or adiponectin gene therapy could improve metabolic health in SKO mice, we performed an oral glucose tolerance test (OGTT), an insulin tolerance test (ITT) and measured *ad lib* fed blood glucose 6 weeks post injection. As shown previously^19^, oral glucose (Fig. 2A) and insulin (Fig. 2B, Supp. Fig. 1B) tolerance tests were impaired in SKO eGFP mice compared to WT mice. Leptin gene therapy did not alter glucose tolerance (Fig. 2A, Supp. Fig. 1C), but moderately improved insulin sensitivity (Fig. 2B, Supp. Fig. 1B, D) and lowered *ad lib* fed blood glucose levels (Fig. 2C) in SKO mice, although neither reached statistical significance.

**Figure 2.**
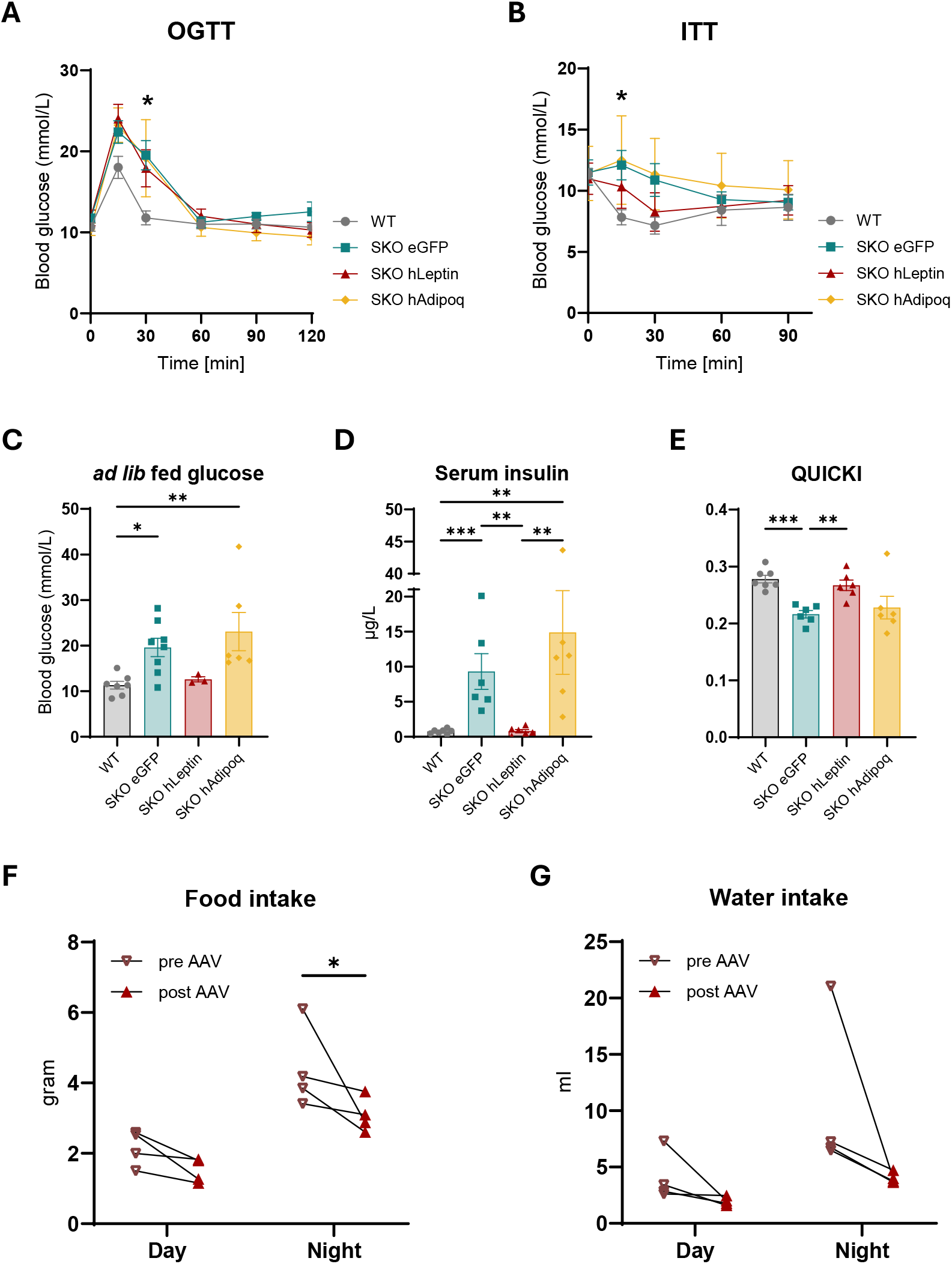
Fasted oral glucose tolerance test (OGTT) (A), fasted insulin tolerance test (ITT) (B) and *ad lib* fed blood glucose levels (C) in WT and SKO mice 6 weeks post gene therapy. Serum insulin levels (D) and QUICKI analysis (E) in WT, SKO eGFP, SKO hLeptin and SKO hAdipoq mice fasted for 5h. Food (F) and water (G) intake in SKO hLeptin mice pre and 6 weeks post AAV injection. Data presented as the mean ±SEM, for A-B n = 7 (WT), 8 (SKO eGFP), 6 (SKO hLeptin), 6 (SKO hAdipoq), for C n = 7 (WT), 8 (SKO eGFP), 3 (SKO hLeptin), 6 (SKO hAdipoq), for D, E n = 7 (WT), 6 (SKO eGFP), 6 (SKO hLeptin), 6 (SKO hAdipoq), for F-G n = 4. For A-B *WT vs SKO eGFP. One-way ANOVA/2way ANOVA, *p≤0.05, **p≤0.01, ***p≤0.001.

SKO eGFP mice had elevated fasted serum insulin levels compared to WT control mice, consistent with observations in CGL2 patients. This fasting hyperinsulinemia was fully normalised in SKO hLeptin mice (Figure 2D). In line with this data, quantitative insulin-sensitivity check index (QUICKI) analysis showed significantly decreased insulin sensitivity in SKO eGFP mice, and this was rescued with leptin gene therapy to values similar to those in WT control mice (Fig. 2E). In contrast, adiponectin treatment failed to elicit changes in these metabolic parameters in both male (Fig. 2A-E) and female SKO mice (Supp. Fig. F-J).

### Effect of leptin gene therapy on food and water intake

Leptin replacement is an established treatment for lipodystrophy, used to improve metabolic abnormalities such as insulin resistance and hyperphagia^20,21^. We compared the effects of leptin gene therapy on food and water intake before and after AAV delivery, each over a 4-day period. Consistent with the clinical features observed in patients with CGL2, we have previously shown that SKO mice exhibit hyperphagia and elevated water consumption compared to WT controls^17^, reflecting polydipsia, a hallmark of metabolic dysregulation commonly seen in diabetic states^22^. Following leptin gene therapy, food intake was significantly reduced and water intake lowered in SKO mice (Fig. 2F, G).

### Leptin but not adiponectin rescues hepatic steatosis in SKO mice

Reduced hepatic steatosis is a significant benefit of recombinant leptin treatment in CGL2 patients. But to date, little is known about the effect of adiponectin treatment in patients with this condition. We therefore assessed the effect of our leptin and adiponectin gene therapy on the livers of SKO mice. As expected, liver weight was significantly higher in SKO eGFP mice compared with WT controls (Fig. 3A). This hepatomegaly was reversed in SKO hLeptin mice (Fig. 3A) but not SKO hAdipoq mice (Fig. 3A, Supp. Fig. 2K). When examining other tissues, the significant reduction in BAT mass observed in SKO eGFP mice was not reversed by leptin nor adiponectin gene therapy, nor was the organomegaly in the pancreas and spleen of SKO eGFP mice rescued in SKO hLeptin or SKO hAdipoq mice (Fig. 3B). We observed increased kidney weights in SKO hLeptin mice, which may merit further investigation (Fig. 3B). When lipid content of livers was examined, we found that the significant increase in hepatic triglycerides in the SKO eGFP mice was entirely normalised in the SKO hLeptin mice 7 weeks after AAV administration (Fig. 3C). Serum ALT levels, a marker of hepatic dysfunction, were modestly elevated in SKO mice which appeared reduced by leptin gene therapy, although neither was statistically significant (Fig. 3D). However, liver lipids in SKO mice were not improved by adiponectin gene therapy and ALT levels of SKO hAdipoq male mice were significantly elevated compared to WT control mice (Fig. 3C, D, Supp. Fig. 2L, M). The substantial reduction of hepatic steatosis in SKO hLeptin versus SKO eGFP and SKO hAdipoq mice was also evident when liver tissue was examined histologically (Fig. 3E, Supp. Fig. 2N). Given the absence of detectable effects with adiponectin treatment, subsequent experiments were conducted in mice receiving leptin gene therapy. Western blotting revealed increased levels of the lipid droplet protein PLIN 2 in SKO eGFP mice which was reduced substantially in SKO hLeptin mice, consistent with changes in intracellular TGs (Fig. 3F). We also examined expression of several inflammatory and fibrosis markers. This revealed elevated levels of *Cola1* and *Col3a1* mRNA in SKO eGFP versus WT mice but these were not normalised by gene therapy in SKO hLeptin mice (Fig. 3G).

**Figure 3.**
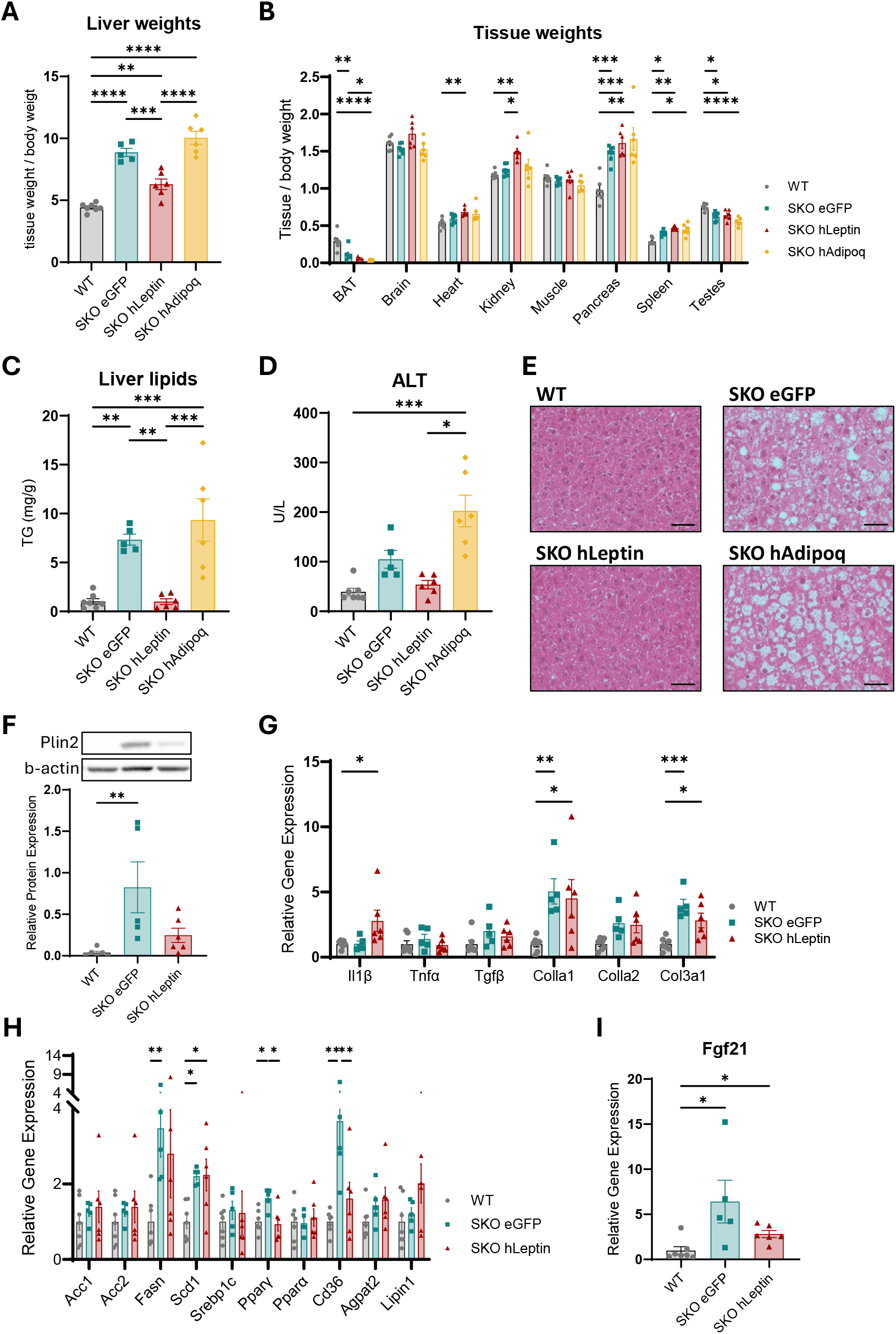
Liver tissue weights (A) and other tissue weights (B) normalised to body weight from WT, SKO eGFP, SKO hLeptin and SKO hAdipoq mice 7 weeks after AAV injection. Liver TG levels (C), ALT serum levels (D) and H&E staining of liver sections (E) from WT, SKO eGFP, SKO hLeptin and SKO hAdipoq mice 7 weeks after AAV injection. Western blot image and quantified relative protein expression of PLIN2 (F). Relative gene expression of fibrosis and inflammatory markers (G), metabolic markers (H) and *Fgf21* (I) in liver of WT, SKO eGFP, SKO hLeptin and SKO hAdipoq mice 7 weeks post AAV injection. Data presented as the mean ±SEM, for A-I n = 7 (WT), 5 (SKO eGFP), 6 (SKO hLeptin), 6 (SKO hAdipoq). One-way ANOVA, *p≤0.05, **p≤0.01, ***p≤0.001, ****p≤0.0001. Scale bar represents 50 µm.

### Leptin normalises expression of *Ppar*γ and *Cd36* in livers of SKO mice

Leptin plays a crucial role in regulating hepatic lipid metabolism by supressing lipogenesis, independently of its effects on food intake^23^. Loss of Seipin leads to increased *de novo* lipogenesis, contributing to hepatic lipid overload and metabolic dysfunctions^24^.

As consistent with previous studies^15,24^ expression of the lipogenic genes *Fasn* and *Scd1* were increased in SKO eGFP mice (Fig. 3H). However, they were not reduced by AAV-leptin administration. The expression of *Srebp1c, Acc1/2, Agpat2* and *Lipin* were not significantly changed in SKO eGFP mice (Fig. 3H). Expression of the key lipogenic transcription factor *Ppar*γ and the fatty acid translocase *Cd36* were significantly increased in SKO eGFP mice but normalised in SKO hLeptin mice (Fig. 3H). We observed increased expression of mRNA encoding *Fgf21* in SKO eGFP mice, consistent with previous studies^25^ and this was also reversed in SKO hLeptin mice (Fig. 3I).

## DISCUSSION

Several preclinical studies have investigated the therapeutic potential of the adipokines leptin^26,27^ and adiponectin^28,29^, or both^30^ delivered *via* gene therapy, in obese mouse models of insulin resistance and fatty liver disease, including diet-induced leptin deficient ob/ob mice.

Here we examine AAV-mediated gene therapy delivering leptin to a preclinical model of CGL, the condition for which daily recombinant leptin injections are currently the most effective approved therapy. Moreover, our study is the first to investigate the therapeutic potential of adiponectin in CGL, arguably the only condition in which affected individuals have dramatically reduced levels of this adipokine. Our data imply that leptin gene therapy could be highly effective in these individuals while we find no evidence for beneficial effects of adiponectin delivery.

Although leptin gene therapy led to marked improvements in liver health, surprisingly, we observed minimal changes in key hepatic metabolic marker genes. Of those examined, only the expression of *Ppar*γ and its downstream target *Cd36* were significantly reduced by leptin gene therapy. It is known that CD36 is upregulated in fatty liver disease, where it facilitates fatty acid uptake and thereby contributes to steatosis^31^ which might suggest a key mechanism by which leptin attenuates hepatic lipid accumulation.

Although human adiponectin was detectable in serum, we did not observe any beneficial effect on glycaemic control and liver health in SKO hAdipoq mice. This contrasts with previous studies reporting beneficial effects of adiponectin in mouse models of insulin resistance^28,29^ including leptin deficiency^9^. Several factors may explain this. Firstly, the levels of serum adiponectin reached following AAV-adiponectin administration were substantially lower than circulating levels of endogenous murine adiponectin we have detected previously in control mice^15,19^, which may have limited its effect. Secondly, we expect that the majority of the adiponectin from the AAV transgene is likely to come from the livers of the SKO mice, particularly given the severe lack of adipose tissue present. It is possible that adiponectin secreted from adipose tissue, as may occur in previous studies of obese mice, is more effective due to secondary modifications and oligomerisation. Thirdly, it may be that the beneficial effects of adiponectin observed in other models requires functional adipose tissue, which is effectively absent in our SKO mice and in CGL patients.

In a previous study using SKO mice^15^, we demonstrated that AAV-mediated gene therapy results in long lasting protein expression from the delivered transgene in liver which persisted for at least five-months post AAV injection. That study and the work described here used the same AAV serotype and CMV promoter. This supports the potential of leptin gene therapy to provide a much longer lasting, sustained treatment for lipodystrophy in the future.

Recent studies, including our own^14,19,30^ highlight the importance of a careful selection of AAV serotypes and tissue specific promoters to ensure that the transgene is efficiently delivered. Building on these findings, future experiments should explore alternative vectors to refine leptin and adiponectin AAV-treatment, potentially identifying novel combinations that lower the systemic dosing, optimise targeting and reduce the likelihood of adverse effects. While AAV8 has shown high transduction efficiency in hepatocytes, alternative serotypes such as AAV1, AAV2 or AAV-DJ -a chimera of type 2/type 8/type 9, along with tissue specific promoters, could potentially offer improved specificity^14^. Notably, these serotypes may show better evasion of immune neutralisation, an essential feature for sustained gene therapy efficacy^32^.

Overall, whilst we found no evidence of beneficial effects of adiponectin delivery, our study defines for the first time the significant potential of leptin gene therapy in a well-recognised model of congenital generalised lipodystrophy. We propose that further refinement and clinical application of this approach could offer a significantly improved, more cost effective, longer lasting and less painful means to deliver this effective treatment for those living with generalised lipodystrophy.

## DATA AVAILABILITY

The datasets generated and/or analysed during the current study are available from the corresponding author on reasonable request.

## ACKNOWLEDGEMENTS

The authors would like to thank the staff at the University of Aberdeen’s Microscopy and Histology Core Facility and the Medical Research Facility for support with animal breeding and maintenance. This work was supported by the MRC MDU Mouse Biochemistry Laboratory (MC_UU_00014/5). Figure 1C “Schematic of experimental procedure” was created with BioRender.com.

## AUTHOR CONTRIBUTIONS

N Sommer: Designed, performed, analysed and interpreted the experiments. Wrote original manuscript, review and editing, funding acquisition.

A Roumane: Performed experiments, reviewed and edited manuscript.

M Tiwari: Performed experiments, reviewed and edited manuscript.

Weiping Han: Resources, reviewed manuscript.

L K Heisler: Resources, reviewed and edited manuscript, project administration.

G D Mcilroy: Resources, reviewed and edited manuscript.

J J Rochford: Project administration, resources, conceptualisation, funding acquisition, supervision, wrote original manuscript, review and editing.

## ETHICS STATEMENT

Animal procedures were approved by the University of Aberdeen Ethics Review Board. The study was conducted in accordance with the local legislation and institutional requirements and performed under the project license PPL PP4951597 approved by the UK Home Office.

