## Supplementary Figure Legends for "AAV-mediated delivery of leptin but not adiponectin improves metabolic health in a mouse model of congenital generalised lipodystrophy"

**Figure S1:** Body weight gain progression in percentage in WT and SKO mice following gene therapy (A). Fold change of ITT (B) and areas under the curve (AUC) using only values above baseline for OGTT (C) and net AUC relative to baseline for ITT measured in WT, SKO eGFP, SKO hLeptin and SKO hAdipoq mice 6 weeks post AAV injection.

Data presented as the mean ±SEM, for A-D n = 7 (WT), 8 (SKO eGFP), 6 (SKO hLeptin), 6 (SKO hAdipoq). For A-B *WT vs SKO eGFP, ‘WT vs SKO hAdipoq, #SKO eGFP vs SKO hLep. One-way ANOVA/2-way ANOVA *p≤0.05, **p≤0.01, ***p≤0.001.

**Figure S2:** Fat mass and fat mass ratio (A-B), lean mass and lean mass ratio (C-D) measured by ECHO-MRI and detection of circulating human adiponectin in serum from AAV injected female SKO mice fasted for 5 hours (E). Fasted oral glucose tolerance test (OGTT) (F), fasted insulin tolerance test (ITT) (G), *ad lib* fed blood glucose levels (H), serum insulin levels (I) and QUICKI analysis (J) in female WT and SKO mice 6 weeks post gene therapy. Liver weights normalised to body weights (K), liver TG levels (L), circulating serum levels of ALT (M) and H&E-stained liver sections (N) of female WT, SKO eGFP and SKO hAdipoq mice 7 weeks post AAV injection.

Data presented as the mean ±SEM, for A-N n = 4 (WT), 6 (SKO eGFP), 7 (SKO hAdipoq). For F-G *WT vs SKO eGFP, ‘WT vs SKO hAdipoq. One-way ANOVA/2-way ANOVA *p≤0.05, **p≤0.01, ***p≤0.001, ****p≤0.0001. Scale bar represents 50 µm.
