## Supplementary figures and images for "AAV-mediated delivery of leptin but not adiponectin improves metabolic health in a mouse model of congenital generalised lipodystrophy"

### Supplementary Figures

#### Supplementary Figure 1:

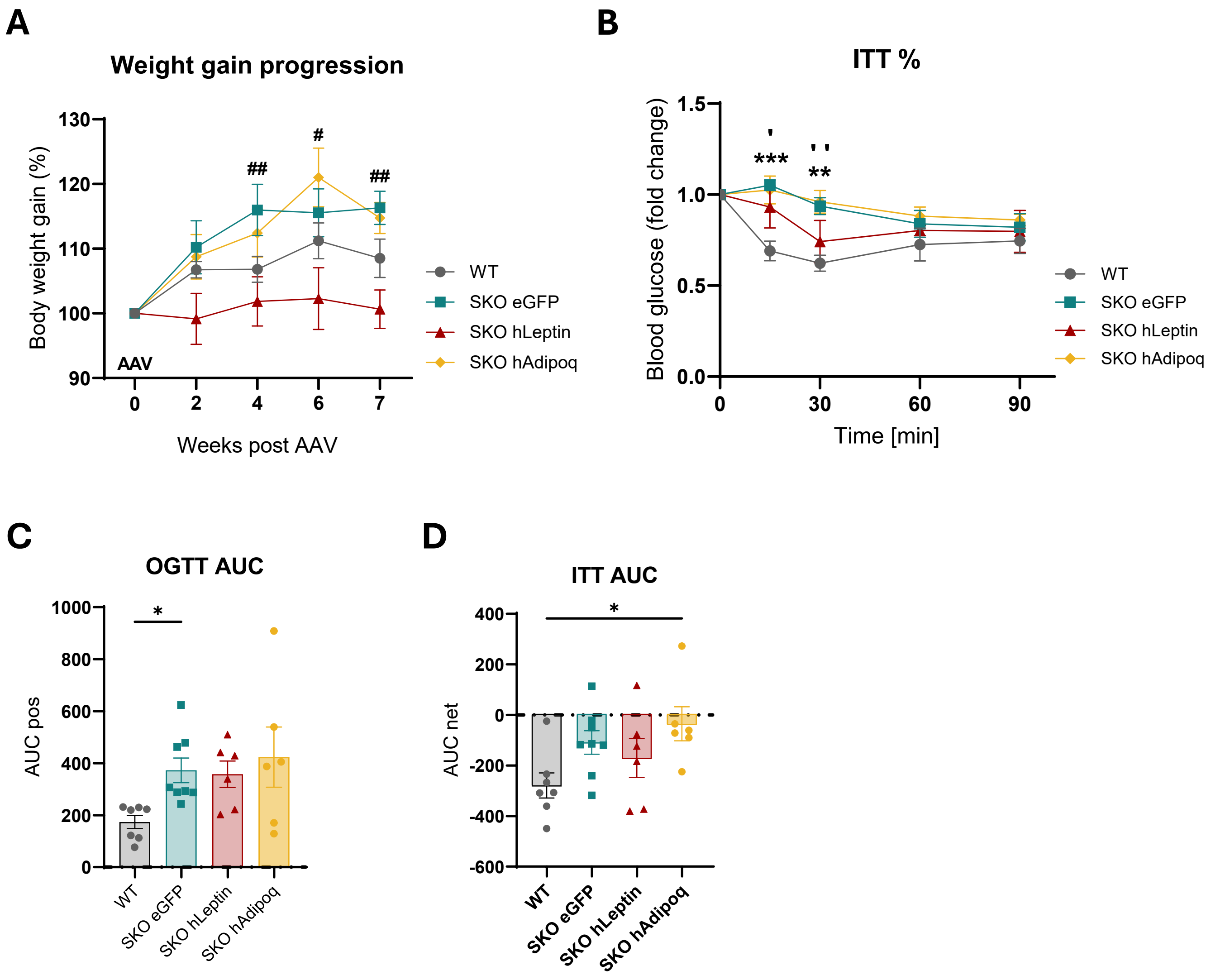

#### Supplementary Figure 2:

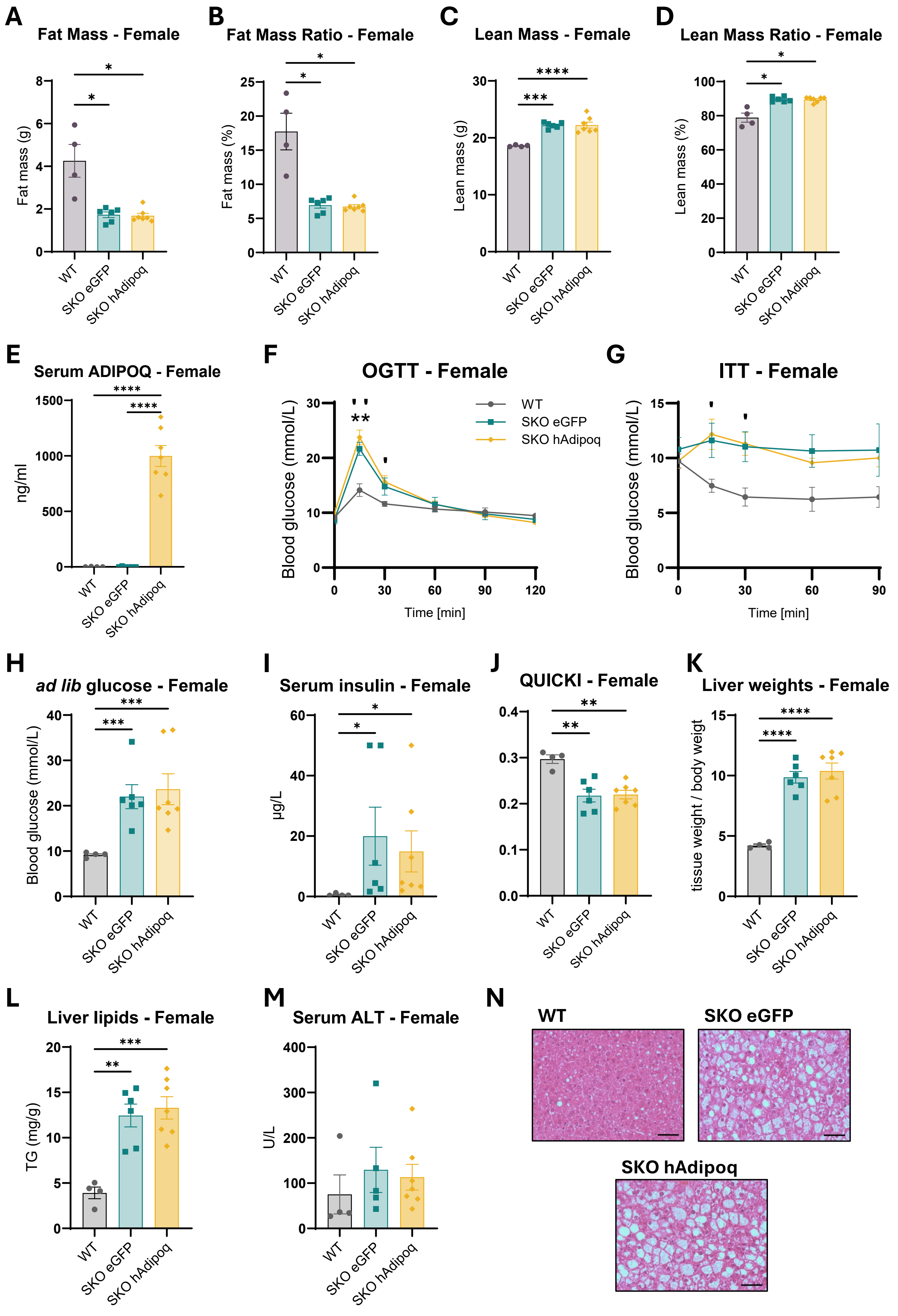
